# Ambroxol displaces α-synuclein from the membrane and inhibits the formation of early protein-lipid coaggregates^†^

**DOI:** 10.1101/2025.08.15.670491

**Authors:** Jesper E. Dreier, Alisdair Stevenson, Elliot Carles, Thomas CT Michaels, Céline Galvagnion

## Abstract

Parkinson’s disease (PD) is a neurological disorder characterized by neuronal loss and the deposition of *α*-synuclein-lipid coaggregates in the brain of patients as well as disruptions in lipid metabolism. Mutations in the gene *GBA*, which encodes the lysosomal glycoprotein Glucocerebrosidase, are together the most important genetic risk factor for PD and have been associated with lysosomal dysfunction, accumulation of pathological *α*-synuclein as well as major changes in both the levels and properties of lipids. Ambroxol, a small molecule chaperone capable of binding and stabilizing Glucocerebrosidase, was found to revert changes in lipid levels and increase in *α*-synuclein levels due to *GBA* mutations potentially via restoring lysosomal function. Here, we show that Ambroxol also has a direct effect on *α*-synuclein-lipid co-aggregation by inhibiting the primary nucleation step in the aggregation process. We find that Ambroxol not only displaces *α*-synuclein from negatively charged membranes but also prevents the formation of early *α*-synuclein-lipid coaggregates during primary nucleation. These results suggest that Ambroxol may have beneficial effects on other synucleinopathies, such as multiple system atrophy and dementia with Lewy Bodies, that are also characterised by the aggregation of *α*-synuclein into amyloid fibrils.

## 1 Introduction

Parkinson’s disease (PD) is the second most common neurodegenerative disease, second only to Alzheimer’s disease and is characterized by the loss of dopaminergic neurons which leads to bradykinesia, tremors and dementia ^1,2^. Another hallmark of PD is the formation of protein-lipid inclusions in the brain called Lewy bodies, whose main protein constituent is *α*-synuclein (*α*S) ^3,4^.

*α*S is a small (14 kDa) intrinsically disordered protein (IDP) that is found at micro-molar concentrations in neuronal synapses ^5^, where its natural function is proposed to be involved in the budding and merging of synaptic vesicles ^6^. In Lewy bodies, however, *α*S is not found in a disordered state, but as rigid amyloid fibrils, where *α*S monomers aggregate through *β*-sheet stacking. Multiple factors have been shown to influence the propensity of *α*S to form amyloid fibrils *in vitro*, such as pH, temperature, metal ions and surfaces such as air-water, polystyrene/water ^7^, membrane/water ^8^ or detergent/water ^9^ interfaces. In particular, the interaction between *α*S and lipid membranes has been shown to play a dual opposite role by being crucial for the proposed biological function of the protein, i.e. synaptic plasticity, while being required for the initiation of the formation of lipid-protein coaggregates ^6,8,10^. The molecular events responsible for the switch from functional to deleterious *α*S-membrane interactions are not yet fully understood. It is now well established that *α*S can bind to membranes made of different lipids, including negatively charged phospholipids, gangliosides and cardiolipin ^11–13^, but is only able to coaggregate with some of these lipids into lipidprotein coaggregates at high protein:lipid ratios ^11,14^. In the case of 1,2-Dimyristoyl-sn-glycero-3-phospho-L-serine (DMPS) membranes, we have recently provided the mechanism by which *α*S co-assembles with DMPS molecules to form lipid-protein coaggregates ^15^. Using tools from chemical kinetics modeling in combination with experimental data, we found that the mechanistic pathway is initiated with a two-step nucleation process at the surface of the membrane followed by an elongation step involving both protein and lipid molecules and thereby consuming lipids from the membrane as the reaction progresses ^15^.

Disruptions in lipid levels have been found to be associated with PD ^16,17^. In particular, mutations in the gene *GBA* encoding the enzyme Glucocerebrosidase (GCase), has been shown to be together the most important genetic risk factor for developing PD and are associated with alterations in the levels and properties of GCase’s substrates glycosylceramide and glucosylsphingosine and more broadly of sphingolipids ^18,19^. *GBA* mutations were found to lead to a decrease in GCase activity and/or protein levels, lysosomal dysfunction and increased levels of lipids as well as total, oligomeric and pathological *α*S in cellular and animal models ^20,21^ of PD and in patients derived samples, including cerebrospinal fluid (CSF), post-mortem brain samples and fibroblasts ^17–19^. Similar findings were observed when GCase activity was reduced upon treatment with small molecule inhibitors such as Conduritol *β* Epoxide (CBE) ^22,23^.

The decreased levels of GCase protein and activity due to *GBA* mutations can be restored by treatment with small molecules, including the small-molecule chaperone Ambroxol (ABX) (see structure in Figure 1a top) ^24–26^. ABX was initially used to treat airway mucus hypersecretion in infants but was found to also act as a molecular chaperone for GCase ^27^. ABX is proposed to bind to the active site of GCase in the endoplasmic reticulum, to stabilise the structure of the protein and to then dissociate from the protein when the ABX/GCase complex reaches lysosomes, thus leaving the protein active ^28,29^. ABX was found to revert the decrease in levels of GCase activity and protein and the increase in the levels of lipids and *α*S due to *GBA* mutations in PD, PD-GBA and/or GD NCSC-DA neurons and fibroblasts ^18,27,30–35^. ABX also led to a decrease in *α*S levels in SH-SY5Y cells and mice overexpressing *α*S ^30,35,36^. Finally, ABX is currently in phase 2 and 3 of clinical trials as a treatment for GBA associated PD as well as in clinical trials as a treatment for Dementia with Lewy Bodies (DLB) ^37–39^.

**Fig. 1.**
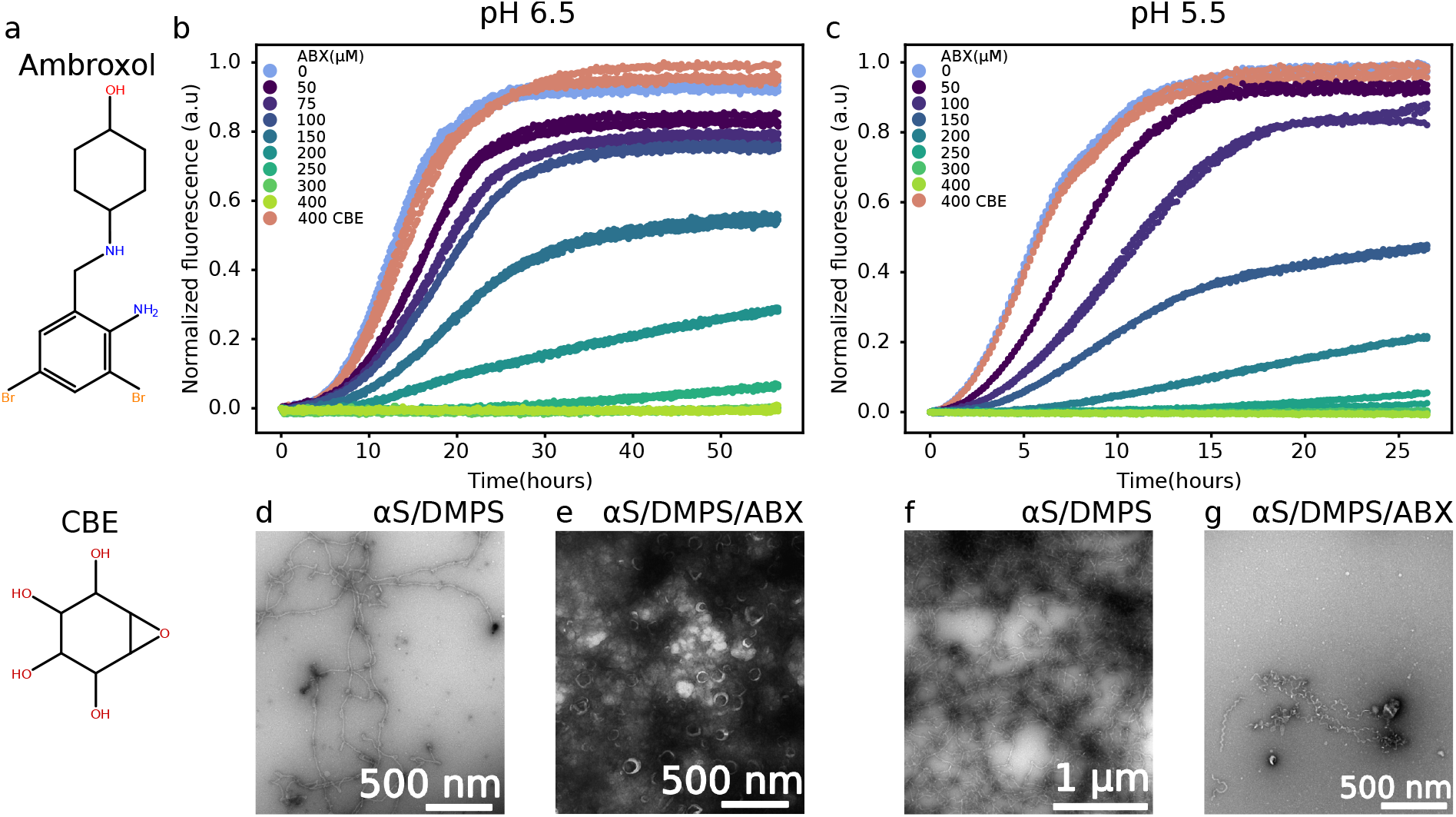
ABX but not CBE inhibits *α*S-DMPS coaggregation into amyloid fibrils at pH 6.5 and 5.5. **(a)** Molecular structure of ABX (top) and CBE (bottom). **(b,c)**. Change in ThT fluorescence for solutions of 50*µ*M *α*S incubated with 200 *µ*M DMPS model membranes in the absence and presence of different ABX or CBE concentrations at pH 6.5 **(b)** and 5.5 **(c)** in non-binding plates under quiescent conditions at 30^*°*^C. d-g. Negative stain EM images of the reaction mixtures 50*µ*M *α*S and 200 *µ*M DMPS model membranes at pH 6.5 **(d-e)** and pH 5.5 **(f,g)** in the absence **(d,f)** and presence **(e,g)** of ABX after a 55 hours **(d,e)** or 27 hours **(f,g)** incubation.

Even though several studies have shown that ABX can decrease the levels of total and/or phosphorylated *α*S in cellular and animal models of PD (including *GBA* mutation and overexpression of *α*S), the mechanism behind this protective effect of ABX against accumulation of *α*S, which could be via an increase in the clearance of *α*S aggregate and/or inhibition of *α*S aggregation, is not yet established.

In the present study, we show that ABX directly prevents *α*S DMPS coaggregate formation and displaces *α*S from the membrane. By using chemical kinetics modeling of lipid-induced protein aggregation with aggregation kinetic data measured for various DMPS:*α*S:ABX ratios, we reveal that ABX inhibits the primary nucleation step in the aggregation pathway of *α*S-DMPS coaggregation. We also find that the inhibitory effect of ABX is specific to this molecule as CBE was found to affect neither *α*S-membrane interactions nor DMPS-*α*S coaggregation into amyloid fibrils.

Our findings show that a direct inhibition of *α*S-lipid fibril formation could be a secondary mechanism of ABX in the treatment of PD in addition to the well characterised GCase chaperone function.

## 2 Results and Discussion

### 2.1 Effect of ABX on *α*S aggregation

To investigate the effect of ABX on the aggregation of *α*S, we incubated *α*S and DMPS model membranes in the absence and presence of different concentrations of ABX under quiescent conditions in non-binding plates (Corning 3881) at pH 6.5 and pH 5.5. We chose to work at pH 6.5 because the process of coaggregation of *α*S and DMPS lipids into mixed protein-lipid fibrils is well characterized by a range of biophysical methods under these conditions ^8,14,^ and the mechanistic pathway of this process has been recently characterized at this pH ^15^. We also studied the effect of ABX on *α*S and DMPS fibrillation at pH 5.5, a value close to that found in the lysosomes.

In the presence of a 1:4 molar ratio of DMPS, *α*S-lipid coaggregates started growing after a lag phase, that was much shorter at pH 5.5 than at pH 6.5 (Figures 1b,c, light blue data points). This observation is in agreement with the faster aggregation of *α*S previously observed under more acidic conditions ^41^. The presence of coaggregates was confirmed by negative stain electron microscopy (EM) (Figure 1d-g). At both pH values, the coaggregates had the same morphology as that of coaggregates formed at pH 6.5 under the same conditions, previously determined using Atomic Force Microscopy ^8^ and cryo-EM ^40^. With increasing ABX concentrations, the rate of aggregation decreased until an ABX:*α*S ratio of 6:1 and ABX:DMPS ratio of 3:2 where no aggregation was observed (Figures 1b,c). Again, this was confirmed with EM where it was only possible to find membranes and no coaggregates in the sample at pH 6.5 (Figure 1e). At pH 5.5, it was possible to also find EM visible coaggregates in the sample with ABX (Figure 1g), but at a much lower abundance than without ABX (Figure 1f). The structures seen in Figure 1g are likely to be aggregated membranes clustering together.

To test if the inhibitory effect observed for ABX was specific to this molecule or generic for small molecules affecting GCase activity/protein levels, we measured the aggregation of *α*S and DMPS under the same conditions as those described so far in the presence of CBE instead of ABX. CBE concentration was set to 400 *µ*M to match that of ABX, where complete inhibition of the DMPS-induced *α*S aggregation was observed. We found that CBE had no effect on *α*S and DMPS coaggregate formation at either pH values (Figures 1c,d), suggesting that ABX specifically inhibits this process.

Having established the inhibitory effect of ABX in the lipid-dependent coaggregation of *α*S, we wanted to investigate if ABX could also inhibit lipid-independent fibrillation of *α*S. We performed aggregation assays in polystyrene microwell plates (Corning 3601), where the first step of *α*S aggregation is proposed to occur at the polystyrene/water interface ^7^. Here, there was no difference in aggregation rates between *α*S alone and with either CBE or ABX, suggesting that the compounds have no effect on the lipid-independent aggregation of *α*S (Figure S1a). Finally, we also investigated whether ABX had any effect on the elongation step of lipid-independent fibril formation, by incubating *α*S monomers with preformed *α*S fibril seeds and ABX at different concentrations. We did not observe changes in aggregation rate at any ABX concentrations in this experiment, further indicating that it is only the lipid-induced coaggregation of *α*S that is affected by ABX (Figure S1b).

### 2.2 Effect of ABX on *α*S-membrane binding

In the light of the observation that ABX specifically inhibits *α*SDMPS co-aggregation into amyloid fibrils, we then set out to investigate whether ABX may interfere with the binding of *α*S to DMPS membranes, as observed for other small molecule inhibitors such as squalamine ^42^ and trodusquemine ^43^. As *α*S forms an alpha-helix upon binding to membranes made of negatively charged lipids, we used circular dichroism (CD) spectroscopy to probe the interaction. The CD signal of *α*S in the presence of an excess of DMPS membranes (*α*S:DMPS ratio of 1:50) displayed the canonical alpha helical signal as previously reported for *α*Slipid complexes, with minima at 208 nm and 222 nm (Figure 2a) ^8,12,44–46^. With increasing ABX concentrations, the CD signal reverted to that of a protein with a random coil conformation, characteristic for IDPs, including *α*S monomer, in solution in the absence of lipid membranes. The return to random coil conformation indicates that the interaction between *α*S and DMPS was disrupted by ABX. Since there were no changes to the CD signal of *α*S with ABX in the absence of DMPS, these results suggest, together with the lipid-independent aggregation experiments (Figure S1), that ABX out competed the *α*S-DMPS interaction and did not directly interact with *α*S. To obtain more quantitative measurements of the affinity of ABX to DMPS membranes, we determined the fraction of bound *α*S to DMPS membranes for each ABX concentration from the CD signal at 222nm and equation 1 and 2^†^. We then fit the fraction bound to a competitive binding model, as previously done for *β*-synuclein and other small molecule inhibitors ^42,47^. The model quantitatively captures the data accurately and, using the known affinity and stoichiometry of *α*S to DMPS membranes, yielded an affinity between DMPS and ABX of *K*_*D*_ = 1 nM and stoichiometry of 1.45 DMPS molecules per ABX molecule in the complex.

**Fig. 2.**
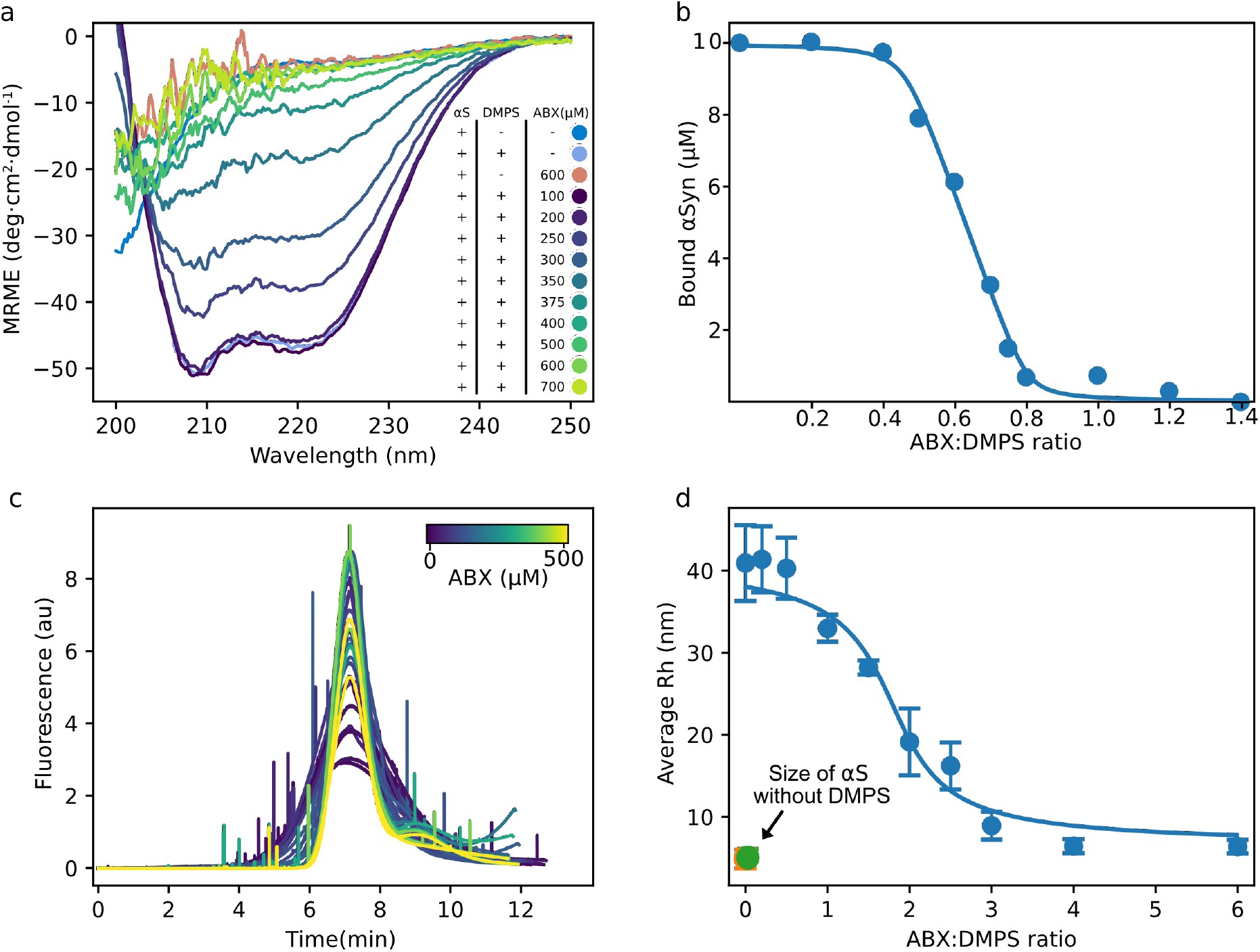
ABX displaces *α*S from DMPS membranes **(a)** Circular dichroism spectra of 10 *µ*M *α*S in the absence and presence of 500 *µ*M DMPS and ABX in phosphate buffer at pH 6.5 and 30^*°*^C. **(b)** Change in the concentration of bound *α*S with increasing ABX concentrations calculated from the MRE at 222 nm using equation 1-2 (see material and methods) and data shown in a. The solid line shows the fit to a competitive binding model between ABX, *α*S and DMPS with the following, previously determined values, for the binding between *α*S and DMPS; *K*_*D*_ = 3.8 µM and stoichiometry of 28.2 DMPS molecules per *α*S monomer. **(c)** Taylor grams from FIDA measurements of 100 nM Alexa 488-labeled *α*S with 30 *µ*M DMPS and increasing concentrations of ABX measured at 30^*°*^C. **(d)** Reported average *R*_*h*_ from the data shown in **(c)**. The orange and green point at 0 ABX:DMPS is *α*S alone and *α*S with 500 *µ*M ABX, respectively. The error bars indicate the standard error between triplicate measurements at each titration point.

To confirm these results using a technique that is independent of the secondary structure change upon binding, we followed the titration of the *α*S-DMPS complex with ABX with flow induced dispersion analysis (FIDA). In this experiment, we measure the hydrodynamic radius (*R*_*h*_) of the fluorescently labeled *α*S from its diffusion rate in the absence and presence of DMPS. Figure 3c shows the raw Taylorgrams of Alexa 488 labeled *α*S with DMPS and increasing ABX concentrations (from dark purple to light yellow). Here, the width of the peak correlates with the diffusion i.e size of the fluorescent molecule. The spikes in the curves indicate a larger, non-diffusive particle in the sample passing the detector. Taylorgrams were converted into diffusion coefficient using the FIDA software (Figure 2d). Figure 2d shows the calculated average size of *α*S in samples with increasing ABX concentrations. In the absence of DMPS and ABX, the calculated size of *α*S was found to be 6 nm, a value similar to that previously determined using microfluidics ^48^, pulsed-field gradient NMR ^49^, fluorescence correlation ^50^ and small-angle X-ray scattering (SAXS) ^51^. The complex size of *α*S with DMPS and no ABX was determined to be approximately 40 nm in radius, a value in the range of what we expected when the DMPS model membranes were produced by extrusion with pore sizes of 100 nm, and is similar to that measured using microfluidics, Dynamic Light Scattering (DLS) and SAXS measurements ^14,48^. With increasing ABX concentrations, the average size of *α*S in the samples was reduced until it reached *∼* 6 nm, corresponding to the size we determined for *α*S without DMPS (orange point in bottom left corner). Interestingly, the ABX:DMPS ratio at which we observed a change in the *α*S:DMPS complex is much higher in the FIDA measurements than it was in the CD measurements. This could suggest that *α*S stays transiently bound to the DMPS model membranes after the alpha helix is lost. The fit of the average *R*_*h*_ of *α*S for increasing ABX:DMPS ratio to the same binding model used for the CD data in Figure 2b gives a *K*_*D*_ of 0.8 *µ*M and a stoichiometry of 0.3 DMPS molecules per ABX molecule in the complex (Figure 2d).

**Fig. 3.**
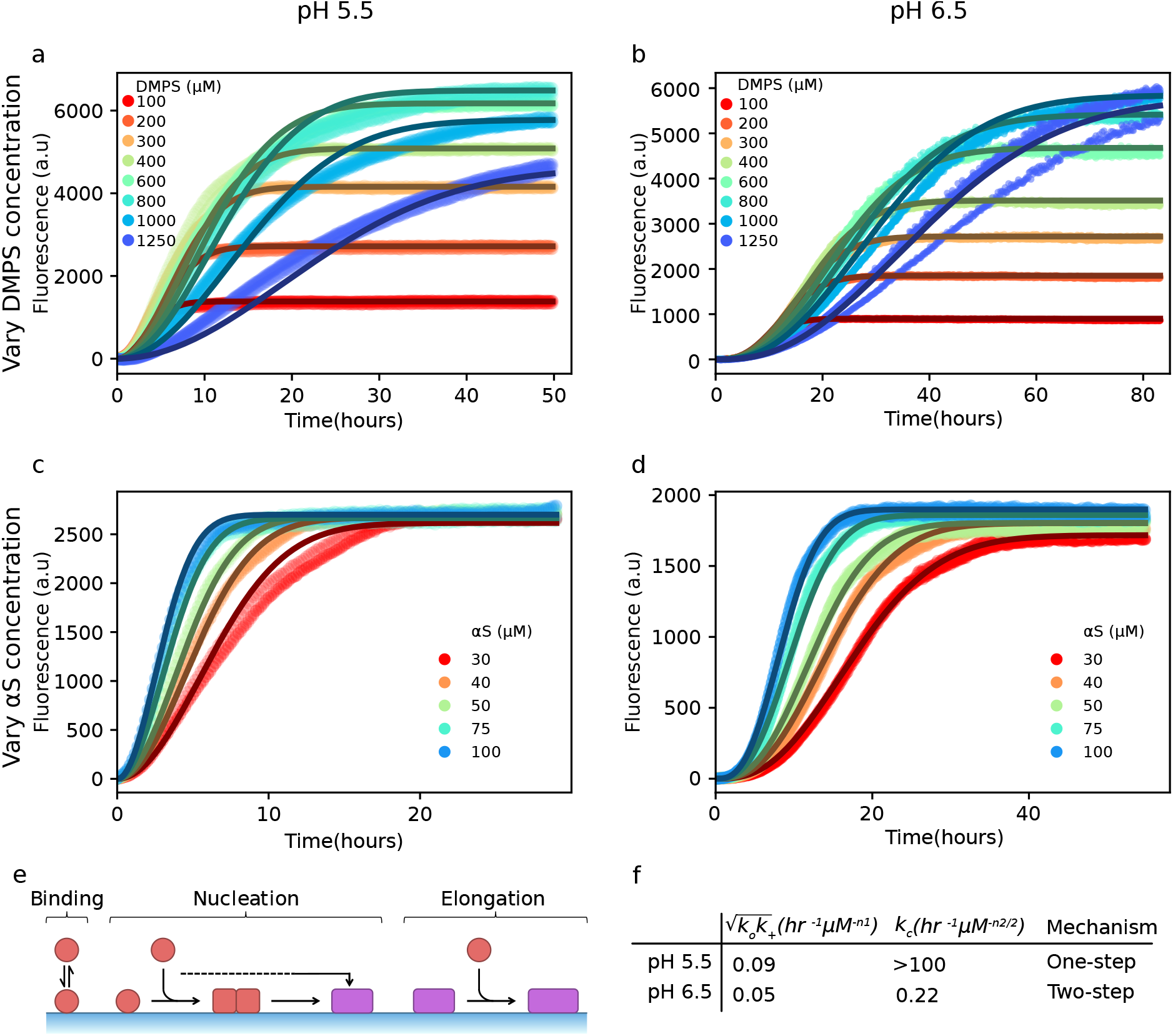
Kinetic analysis reveals the mechanistic difference in *α*S-DMPS coaggregation kinetics between pH 5.5 and pH 6.5 in the absence of ABX. **(a-d)** ThT fluorescence curves from experiments with either a constant concentration of *α*S (50*µ*M) and varying DMPS concentrations (a,b), or a constant concentration of DMPS (200*µ*M) and varying *α*S concentration (c,d). The experiments were performed at both pH 5.5 (a,c) and pH 6.5 (b,d). The aggregation data (scatter points) were globally fit to the integrated rate law equation 5^†^ (solid lines) to determine the values for the different aggregation rate parameters in the absence of ABX. Details on the fitting procedure can be found in the Supplementary Figure 3 and parameters in Supplementary Table 1. **(e,f)** Schematic representation of the microscopic mechanisms in lipid-induced amyloid formation (e) and overview of the rate parameters extracted from fitting the experimental kinetic data (f).

To further investigate the interaction between ABX and DMPS, we performed fluorescence polarization (fp) measurements of DMPS model membranes with the fluorophore 1,6-Diphenyl1,3,5-hexatriene (DPH) and increasing ABX concentrations. The polarization of DPH depends on the fluidity of the DMPS membrane as DPH is buried in the lipid bilayer ^52^. In the absence of ABX, the measured fp was approximately 350, a value characteristic of a membrane in the gel phase, as expected for DMPS at temperatures below the melting temperature of approximately 45^*°*^C (Figure S2). In the presence of increasing ABX concentrations, the measured fp decreased to around 150, a value characteristic of a membrane in the fluid phase. These results indicate that ABX binds to the DMPS membrane and that ABX:DMPS binding leads to the melting of the membrane as observed for *α*S and other small molecules known to displace *α*S from the membrane, such as squalamine ^42^.

Altogether, these results show that ABX displaces the protein from DMPS membranes via a competitive binding mechanism. We next used kinetic analysis to uncover the microscopic steps in the process of DMPS:*α*S coaggregation that ABX is affecting.

### 2.3 Kinetic analysis of lipid-induced *α*S aggregation in the absence of ABX

We first established a baseline kinetic model for the uninhibited system. This kinetic analysis of aggregation in the absence of ABX is essential to later resolve the specific aggregation mechanisms perturbed by ABX. Aggregation dynamics were monitored via ThT fluorescence under two main experimental series: (i) varying initial DMPS concentrations at constant *α*S concentration (Figure 3a,b), and (ii) varying *α*S concentrations at fixed DMPS levels (Figure 3c,d), each conducted at pH 5.5 (Figure 3a,c) and pH 6.5 (Figure 3b,d). Our modeling approach builds on a previously developed chemical kinetics framework for lipid-induced *α*S aggregation ^15^. In this model, the aggregation kinetics are governed by a two-step nucleation process, with oligomerization (with rate constant *k*_*o*_) and conversion (with rate constant *k*_*c*_), followed by coaggregate growth (with rate constant *k*_+_) (Figure 3e). Across all conditions, the experimental data show excellent agreement with the theoretical predictions from our integrated rate laws (solid lines in Figure 3a-d) over a broad range of protein and lipid concentrations, with explicit mathematical details of our model described in the Methods section in the Supplementary Information.

Through globally fitting our model to the experimental data, we can quantify the associated aggregation rate constants (Figure 3f). At pH 6.5, the values for these rate constants confirm a two-step nucleation mechanism in which both oligomer formation and conversion are kinetically relevant. In contrast, the pH 5.5 data yield a markedly higher value of *k*_*c*_, consistent with a regime where primary nucleation becomes effectively one-step. The shift from two-step to one-step nucleation with decreasing pH is especially clear when examining the initial time dependence of the aggregate mass concentration, *M*(*t*). According to our integrated rate laws, one-step nucleation yields a quadratic scaling at early times, *M*(*t*)*∼ t*^2^. In contrast, a two-step mechanism gives rise to higher-order time dependence, typically *M*(*t*)*∼ t*^*n*^ with 2 *< n <* 3, reflecting the additional oligomerization step before conversion ^15^. Analysis of the early-time data yields a scaling close to *M*(*t*)*∼ t*^3^ at pH 6.5, which shifts to a scaling close to *M*(*t*)*∼ t*^2^ at pH 5.5 (Figure S3).

### 2.4 Mechanism of inhibition of lipid-induced *α*S aggregation by ABX

Next, we sought to understand how ABX inhibits the *α*S-DMPS coaggregation into amyloid fibrils. To this end, we employed the same two-step kinetic model used to describe the uninhibited system and, by leveraging the parameters previously extracted from the uninhibited kinetics (Figure 3f), we systematically perturbed individual microscopic steps in the model to capture the effect of ABX on aggregation. Kinetic traces reveal the impact of varying concentrations of ABX on aggregation kinetics at pH 5.5 and 6.5 (Figure 4a and b, respectively). The decrease in plateau fluorescence recorded in Figure 4a,b indicates that the yield of coaggregates decreases with increasing ABX concentrations, assuming a correlation with aggregated protein mass and ThT fluorescence as previously confirmed experimentally ^8^. Given that the yield of amyloid fibril formation is limited by available lipid under our experimental conditions ^15^, and that ABX was found to displace *α*S from DMPS model membranes (Figure 2), we initially considered whether ABX inhibition could be explained solely by a reduction in effective lipid concentration, i.e. by only preventing the binding of *α*S to DMPS membranes. However, model fitting in which only lipid availability was reduced failed to capture the full kinetic profiles (Figure S4), suggesting that ABX is also directly inhibiting a microscopic reaction in the aggregation pathway.

**Fig. 4.**
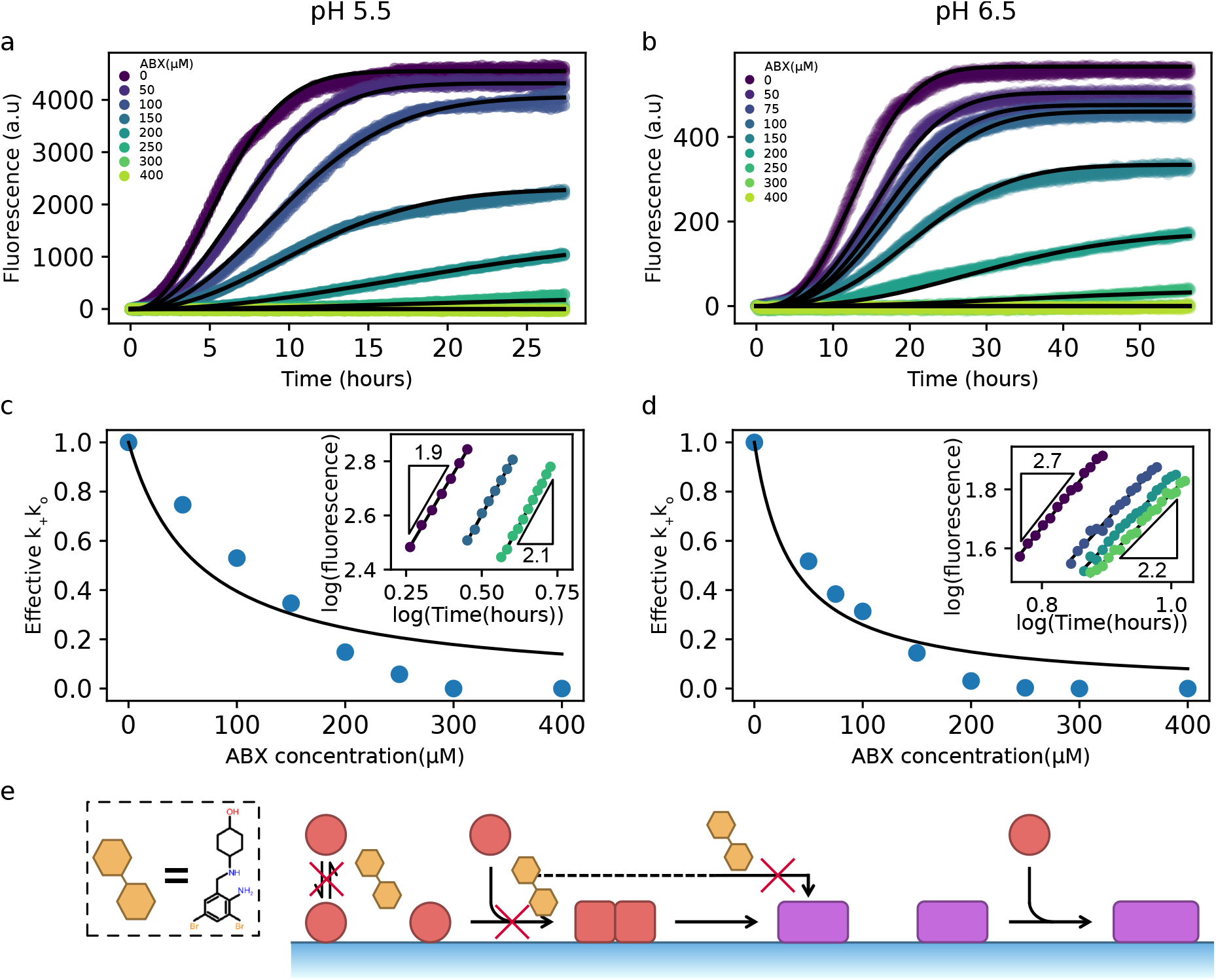
Identification of microscopic mechanism through which ABX affects *α*S-lipid co-aggregation. **(a,b)** The experimental ThT profiles (in triplicates) at different ABX concentrations and at pH 5.5. (a) or pH 6.5 (b) are compatible with a our integrated rate law when the rate parameter of oligomer formation, *k*_*o*_, is reduced in a ABX-concentration dependent manner. **(c,d)** Effective microscopic rate constant *k*_*o*_ as a function of ABX concentration, reported relative to the values in the absence of ABX. The black line represents a fit of the prediction from the relationship 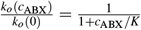, where *c*_ABX_ is the ABX concentration. Insets: Double-logarithmic plots of ThT curves at early times show how ABX affects the early-time scaling *M*(*t*)*∼ t*^*n*^. The slope of the lines corresponds to the exponent *n*. A value *n* =2 indicates a one-step nucleation mechanism, whereas a value *n* =3 indicates a two-step nucleation mechanism. All fitting parameters can be found in Supplementary Table 1. **(e)** Schematic representation of the proposed inhibition mechanism by ABX.

To identify the inhibited step, we therefore perturbed specific microscopic rate constants in our kinetic model in response to changing ABX concentration. This analysis indicates that our experimental data are consistent with a scenario where only the oligomer formation rate *k*_+_*k*_*o*_ was reduced in an ABX-dependent manner, while *k*_*c*_ was held fixed (Figure 4c,d). This strategy yielded excellent agreement with the data across a wide range of ABX concentrations and at both pH values. The extracted *k*_+_*k*_*o*_ values exhibit a monotonic decrease with increasing ABX concentrations (Figure 4c,d), which can be modeled using a simple relationship of the form 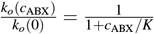, where *c*_ABX_ is the ABX concentration. This analysis yields a characteristic inhibition constant *K ≈* 30 *µM* at both pH 5.5 and 6.5.

Furthermore, the proposed inhibition of oligomer formation as the mechanism of action of ABX is consistent with the changes in the early-time curvature of the kinetic traces as the ABX concentration increases (insets of Figure 4c,d). Specifically, early-time log-log plots of aggregate mass versus time at pH 6.5 show a reduction in the effective scaling exponent from *∼*3 to *∼*2 as ABX concentration increases (inset of Figure 4d). A *t*^3^ scaling is typically indicative of a two-step nucleation process with a rate-limiting oligomer conversion step, while a *t*^2^ scaling implies that the oligomer conversion process is no longer rate-limiting and that, therefore, the oligomer formation process is rate limiting ^53^. The shift towards a quadratic scaling indicates that the presence of ABX slows down the overall aggregation rate by inhibiting oligomer formation, making the conversion step less rate-limiting. At pH 5.5, where *k*_*c*_ is already high and conversion is not rate-limiting, the early-time kinetics remain quadratic, consistent with a picture where ABX targets the oligomer formation step (inset of Figure 4c). Taken together, these results are compatible with a mechanism whereby ABX inhibits *α*S-DMPS coaggregation into amyloid fibrils by interfering with the formation of early oligomeric nuclei (Figure 4e).

## Conclusions

In this study, we show that ABX, a small molecule known to stabilise GCase and restore lysosomal function associated with GBA mutations, can act directly on *α*S by inhibiting its co-assembly with lipid molecules into amyloid fibrils. Using kinetic analyzes of DMPS-induced aggregation curves of *α*S, we discovered that ABX specifically inhibits the primary nucleation step of the process of lipid-protein coaggregation and decreases the rate of formation of early oligomers. The inhibitory effect of ABX was specific to the small molecule as we did not observe an effect of CBE, a small molecule inhibitor of GCase, on DMPS-induced *α*S aggregation. Moreover, ABX specifically inhibits the process of DMPS-*α*S coaggregation into amyloid fibrils as we did not observe any effect of the small molecule chaperone on *α*S aggregation in the absence of lipids, i.e. in polystyrene plates or in the presence of seeds in non-binding plates under quiescent conditions.

In light of the fact that ABX inhibits specifically DMPS-induced *α*S aggregation, we investigated the effect of ABX in the interaction between *α*S and DMPS model membranes using both CD spectroscopy, a spectroscopic method dependent on secondary structure changes upon binding, and taylor dispersion analysis, a diffusion based method independent of secondary structure changes upon binding. The results from the titrations of ABX into the DMPS:*α*S system show that ABX displaces *α*S from the lipid model membranes. Measurements of the fluidity of the DMPS model membranes revealed that increasing ABX concentrations also increased the fluidity of the model membranes (Figure S2). Since ABX has no effect on *α*S aggregation in the absence DMPS (Figure S1), these binding data suggest that ABX does not interact directly with *α*S but displaces the protein from the DMPS membranes via a competitive binding.

Given that pH varies through the lysosomal maturation where ABX performs its chaperonal activity on GCase ^27,28^, and has been shown to change *α*S aggregation rate, we investigated the effect of ABX at both pH 6.5 and 5.5. Aggregation occurred faster at lower pH, as has been shown in previous studies ^41,54^. The kinetic analyses of the aggregation curves at pH 6.5 and 5.5 show that this acceleration is mainly due to an increase in the conversion step. Even under these faster aggregating conditions, ABX still inhibited the DMPS-induced aggregation of *α*S at pH 5.5.

Altogether, these results highlight a dual role for ABX as a small molecule capable of both stabilizing GCase and inhibiting *α*S:lipid coaggregation. ABX is a promising molecule for the treatment of PD and is currently in phase 2 and 3 of clinical trials as a treatment for GBA associated PD as well as in clinical trials as a treatment for dementia with Lewy bodies (DLB) ^37–39^. Our study suggests that ABX may prevent *α*S aggregation in different synucleinopathies including PD, DLB and Multiple System Atrophy.

## Author contributions

Designed research: JED, AS, TCTM, CG; Performed research: JED, AS, EC; Analyzed data: JED, AS, TCTM, CG; wrote the paper: JED, AS, TCM, CG.

## Data availability

Data for this article, including raw experimental data (ThT fluorescence measurements) are available at Zenodo.

## Acknowledgements

This work was supported by ETH Zurich (A.S., T.C.T.M.), the Swiss National Science Foundation (Grant Number SNS 219703) (T.C.T.M.) and the Novo Nordisk Foundation (Grant number: NNF20OC0059417, J.E.D, E.C., C.G. We acknowledge the Core Facility for Integrated Microscopy, Faculty of Health and Medical Sciences, University of Copenhagen.

