## Supplementary Information for "Ambroxol displaces α-synuclein from the membrane and inhibits the formation of early protein-lipid coaggregates^†^"

### Supplemental information for: Ambroxol displaces $\alpha$-synuclein from the membrane and inhibits the formation of early protein-lipid coaggregates

Jesper E. Dreier,[†] Alisdair Stevenson,[‡,¶] Elliot Carles,[†] Thomas CT Michaels,[*,‡,¶] and Céline Galvagnion[*,†]

†*Department of Drug Design and Pharmacology, Faculty of Health and Medical Sciences, University of Copenhagen, 2100 Copenhagen, Denmark.*

‡*Department of Biology, Institute of Biochemistry, ETH Zurich Otto Stern Weg 3 8093 Zurich Switzerland.*

¶*Bringing Materials to Life (BML) Initiative, ETH Zurich, Switzerland*

### Materials and methods

#### Protein purification

Wild type, N-terminally acetylated $\alpha$S was expressed and purified as previously described.[?] Briefly, $\alpha$S and the pNatB acetylation complex were co-expressed in e.coli BL21 pLysS cells. The cells were lysed with osmotic shock buffer, and the protein purified by heat precipitation, ion exchange chromatography and finally size exclusion chromatography. The protein concentration was determined by measuring absorbance at 280 nm. Subsequently, the protein was aliquoted, flash frozen with liquid nitrogen and finally stored at -80 °C

#### DMPS model membrane formation

DMPS was purchased from Avanti Lipids as a powder. The powder was solubilized in a 9:1 chloroform:methanol mixture and aliquoted. DMPS stock solution was added to a round bottom flask, where the solvent was evaporated with nitrogen gas to form a lipid film. The lipid film was dehydrated under vacuum in a desiccator for 1 hour or O/N. Next, the lipid film was resuspended in 20 mM phosphate buffer pH 6.5 for 1.5 hours at 50 °C and finally extruded through polycarbonate membranes with 100 nm pore diameter (Avanti Polar Lipids, Inc) to produce small unilamellar vesicles. The size distribution was confirmed by dynamic light scattering measurements.

#### Aggregation Kinetics

All aggregation experiments were performed with 50 $\mu$M N-terminally acetylated $\alpha$S, unless stated otherwise, 50 $\mu$M ThT and measured on a BMG Labtech clariostar microplate reader under quiescent conditions and at 30 °C. The experiments where carried out in Corning 96 well black polystyrene plates with a non-binding surface. For experiments where the aggregation was induced by a polystyrene, the plates had a high-binding surface. The lipid induced aggregation assays were performed in the presence of 200 $\mu$M DMPS SUVs. Experiments were performed in either 20 mM phosphate buffer pH 6.5, or 10 mM acetic acid – acetate buffer, pH 5.5.

#### Negative stain transmission electron microscopy

Once samples had reached a plateau of aggregation, an aliquot was taken and placed on a carbon grid. Then 2 % uranyl acetate was added to the grid, incubated for 30 seconds and subsequently dried. Finally, the grid was washed with $dH_2O$. Images of the grids were acquired on a Philips CM100 transmission electron microscope at the Core facility for integrated microscopy at the Faculty of Health and Medical Sciences, University of Copenhagen.

#### CD spectroscopy

CD spectroscopy was performed on a Jasco J-1500 spectrometer. All spectra were recorded at 30 °C in a quartz cuvette with a 1 mm path length. Samples contained 10 $\mu$M N-ace-$\alpha$S in 20 $\mu$M phosphate buffer pH 6.5, and indicated concentrations of DMPS, ABX and CBE. Scanning speed: 50 nm/min, bandwidth:1 nm, data pitch 0.1 nm, D.I.T: 1 sec. The reported CD signal is the average of 5 accumulations and calculated as the mean residue molar ellipticity (MRME). The CD signal can be described as:

$$CD_{measured} = CD_{free}x_{free} + CD_{bound}x_{bound} \tag{1}$$

where $CD_{measured}$ is the observed CD signal, $CD_{free}$ is the CD signal of free $\alpha$S, $x_{free}$ is the concentration of free $\alpha$S, $CD_{bound}$ is the signal of bound $\alpha$S and $x_{bound}$ is the concentration of $\alpha$S. If we assume that $x_{free} + x_{bound} = 1$ and that $\alpha$S in the absence and presence of DMPS under saturating conditions has the signals $CD_{free}$ and $CD_{bound}$ respectively, (1) can be rewritten as:

$$[\alpha S_{\text{Bound}}] = \frac{CD_{\text{Obs}} - CD_{\text{Bound}}}{CD_{\text{Free}} - CD_{\text{Bound}}}[\alpha S_{\text{total}}] \tag{2}$$

where $CD_{\text{Obs}}$ is the measured CD signal, $CD_{\text{Bound}}$ is the CD signal of fully bound $\alpha$S and $CD_{\text{Free}}$ is the CD signal of $\alpha$S without DMPS at 222 nm.

To quantify the affinity between ABX and DMPS, the competitive binding model described by Brown et. al and Perni et. al[?,?] was used.

#### Fluorescence polarization

From a stock 96% ethanol, 1,6-diphenyl-1,3,5-hexatriene (DPH) were incubated with DMPS at a 1:300 DPH:DMPS ratio at 45 °C for 30 minutes in a 20 $\mu$M phosphate buffer pH 6.5. Samples with DPH-DMPS model membranes were prepared in a Corning 96-well black polystyrene plate with non-binding surface at 500 $\mu$M DMPS with concentrations of ABX between 0 and 500 $\mu$M. The measurements were performed at temperatures around the melting point of DMPS (35-40 °C) and recorded in a BMG Labtech clariostar plate reader using top optic with an excitation at 360 nm and emission at 450 nm.

#### FIDA

FIDA experiments were performed on the Fida 1 instrument (Fida Biosystems, Denmark) in the standard 70 $\mu$m capillary. 100 nM Alexa-488 labeled $\alpha$S A140C was mixed with 30 $\mu$M DMPS model membranes and varying ABX concentrations as indicated on Fig. **??** in 20mM phosphate buffer, pH 6.5. The data was recorded at 30°C. The FIDA experiments were run with the parameters stated in table 1. The raw Taylorgrams were fitted and the $R_h$ was extracted using the FIDA software[?] using the following equations. First, The apparent diffusion coefficient is calculated as:

$$D_{app} = \frac{a^2}{24\sigma^2} t_R \tag{3}$$

where $Dapp$ is apparent fluorescence, a is inner radius of the capillary, $\sigma^2$ is the variance of the peak and $t_R$ is the residence time.

Next, using the Stoke-Einstein equation, the apparent $R_h$ can be calculated:

$$R_{happ} = \frac{k_b T}{6\pi\eta D_{app}} \tag{4}$$

where $k_b$ is Boltzmann constant, T is the temperature and $\eta$ is the viscosity.

Table S1: FIDA parameters

| Step | Pressure (mPa) | Duration (seconds) | Solution |
|---|---|---|---|
| 1. Wash | 3000 | 60 | NaOH (1M) |
| 2. Equilibration | 3000 | 60 | Buffer |
| 3. prePlug | 200 | 30 | analyte |
| 4. Plug | 200 | 10 | indictaor |
| 5. Run | 50 | 700 | analyte |

#### Aggregation kinetics analysis and effect of inhibitor

From our previous analyses,[?] the generation of lipid-protein coaggregate mass, $M(t)$, when two-step primary nucleation events occur, is described by the following integrated rate law:[?]

$$\frac{M(t)}{m_{\text{tot}}} = \begin{cases} \frac{r\,A}{\alpha}\left(1 - e^{-F(t)}\right) & \text{vary lipid concentration} \\ A\left(1 - e^{-F(t)}\right) & \text{vary protein concentration} \end{cases} \tag{5}$$

where the explicit form of the function $F(t)$ is given by:

$$F(t) = \frac{\lambda^2}{k_c^2}\left(1 - k_c t + \frac{1}{2}(k_c t)^2 - e^{-k_c t}\right) \tag{6}$$

with

$$\lambda = \sqrt{\frac{2k_+ k_o \theta_0^{n_1+1}\left(m_{\text{tot}} - L_{\text{tot}}\frac{\theta_0}{\beta}\right)^{n_2}}{L_{\text{tot}}\,A}}\,. \tag{7}$$

Here,

$$A = \begin{cases} 1\,, & r < \alpha \\ \frac{\alpha}{r}\,, & r \geq \alpha\,, \end{cases} \tag{8}$$

where $r = L_{\text{tot}}/m_{\text{tot}}$ denotes the lipid-to-monomer ratio, $m_{\text{tot}}$ is the initial concentration of protein monomers, $L_{\text{tot}}$ is the lipid concentration, $\alpha$ is the stoichiometry of lipids bound to a protein in fibril form, $\beta$ is the stoichiometry of lipids bound to a protein in monomeric form,[?] and $\theta_0$ is the surface coverage of protein monomers at the early stage of the reaction:

$$\theta_0 = \frac{m_{\text{tot}} + K_D + \frac{L_{\text{tot}}}{\beta} - \sqrt{\left(m_{\text{tot}} + K_D + \frac{L_{\text{tot}}}{\beta}\right)^2 - \frac{4m_{\text{tot}}L_{\text{tot}}}{\beta}}}{2L_{\text{tot}}/\beta}, \tag{9}$$

where $K_D$ is the equilibrium dissociation constant for protein-lipid binding. This integrated rate law accounts for combinations of the rate parameters, including oligomer formation ($k_o$), oligomer conversion ($k_c$), and fibril elongation ($k_+$), that define much of the macroscopic aggregation behavior. To capture the monomer or lipid dependence of these aggregation steps, we use reaction orders, where $n_1$ is the variable reaction order accounting for the dependence of the primary nucleation mechanism on bound protein monomers and $n_2$ is the variable reaction order describing the dependence of nucleation on free protein monomers.

With this integrated rate law, the effect of an inhibitor on the lipid-induced aggregation kinetics can be captured by altering the rate parameters $k_o$, $k_c$ and $k_+$ in the integrated rate laws described in (5). Perturbing the different microscopic rate constants results in characteristic changes in the kinetic profile that, when compared to experimental data, can be used to identify the mechanism of action of an inhibitor without explicitly describing the microscopic interaction between the inhibitor with proteins and lipids.

### Supplementary Table

Table S2: Fitting parameters

| Parameter | pH 5.5 Vary lipid | pH 5.5 Vary protein | pH 6.5 Vary lipid | pH 6.5 Vary protein | pH 5.5 Inhibited | pH 6.5 Inhibited |
|---|---|---|---|---|---|---|
| $k_c$ $(hr^{-1}\mu M^{-n_1})$ | 100 | 100 | 0.22 | 0.24 | 100 | 0.12 |
| $\sqrt{k_+ k_o}$ $(hr^{-1}\mu M^{-n_2/2})$ | 0.08 | 0.1 | 0.08 | 0.012 | Varied, see fig. 4 | Varied, see fig. 4 |
| $n_1$ | 0 | 0 | 0 | 0 | 0 | 0 |
| $n_2$ | 1.1 | 1.1 | 0.8 | 1.3 | 1.1 | 0.8 |
| $\alpha$ | 10 | 10 | 14 | 14 | 10 | 14 |

### Supplementary Figures

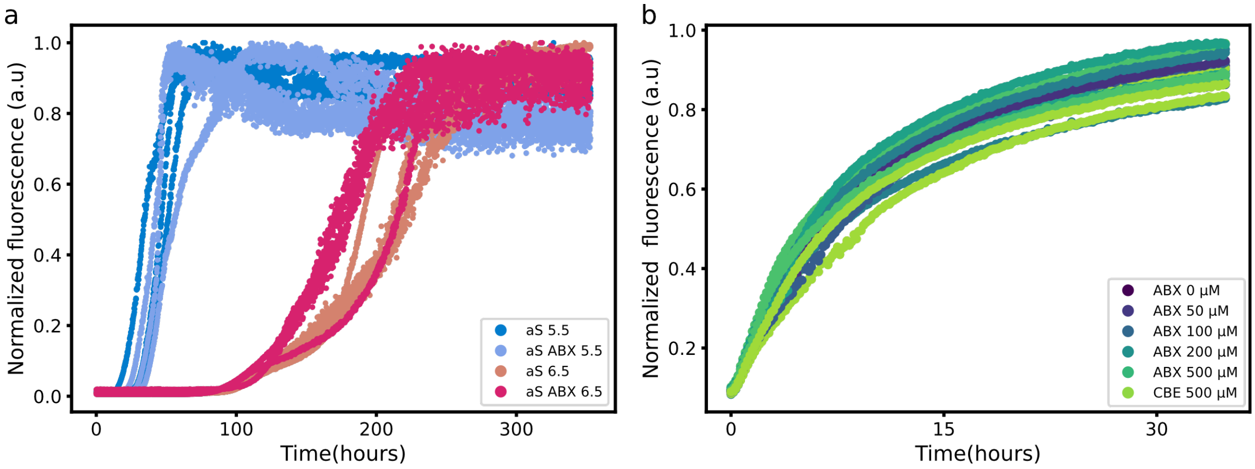


Figure S1: ThT aggregation experiments without lipids are not affected by ABX. **(a)** Polystyrene induced $\alpha$S aggregation performed at pH 5.5 or 6.5, 30°Cin a polystyrene (high-binding plate) with 50 µM $\alpha$S, with or without 400µM ABX, suggests no influence of ABX on the aggregation rate. **(b)** Seeded aggregation with 2 µM preformed $\alpha$S seeds, 50 µM monomeric $\alpha$S and indicated concentrations of ABX at pH 6.5

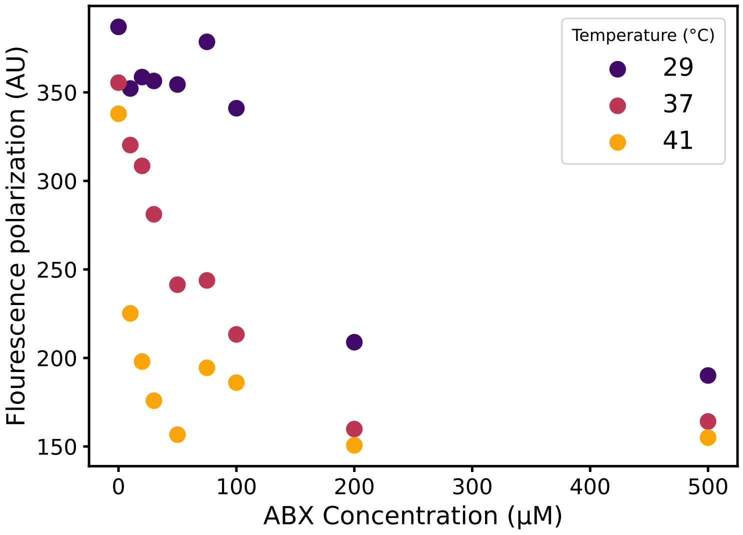


Figure S2: ABX increases the fluidity of DMPS membranes. The figure shows the fluorescence polarization of DPH in DMPS vesicles plotted against ABX concentration. A high value indicates a more rigid lipid bilayer. At increasing ABX concentrations, the polarization drops, indicating a phase transition in the bilayer. At elevated temperatures, the phase transition occurs at lower ABX concentrations. The reported values are the mean of duplicate values.

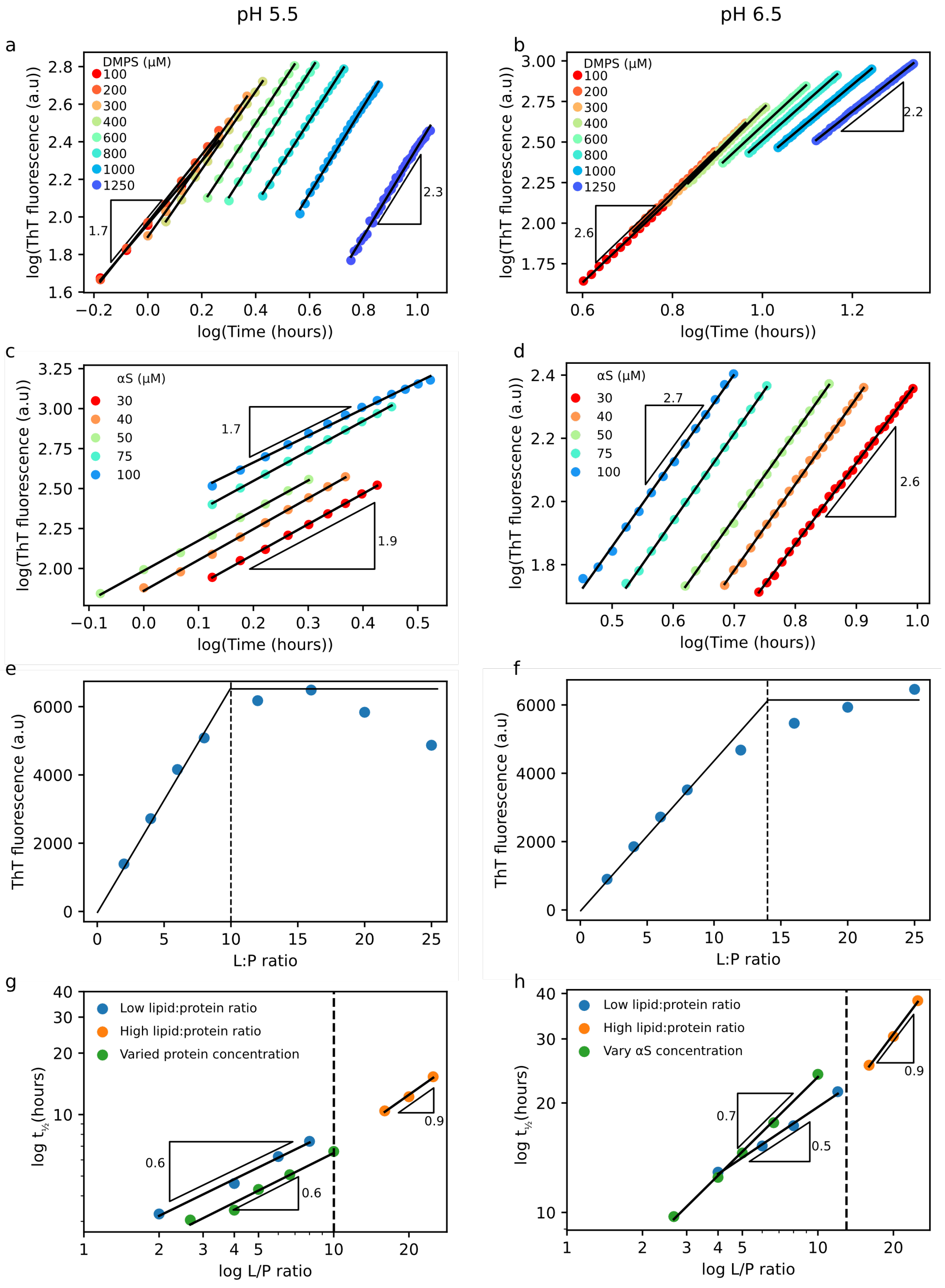


Figure S3: Aggregation parameters extracted for fitting the data. **(a-d)** Early time behavior of the vary lipid **(ab)** and vary protein **(cd)** kinetic curves from fig3 where the logarithm of ThT fluorescence is plotted against the logarithm of the time. The triangles represent the slopes. **(ef)** The ThT fluorescence of each L:P ratio of vary lipid samples at the plateau at the end of the experiment plotted against the L:P ratio. The black lines are the theoretically predicted steady-state behaviour. The stippled line represent the $\alpha$ value which is where the regime changes from being lipid-limited to being protein-limited. **(gh)**. Logarithm of the $t_{\frac{1}{2}}$ of the aggregation curves shown in Fig. 3 plotted against the logarithm of the L:P ratio. The stippled line represents the $\alpha$ value. The triangles visualize the slopes of the relationship.

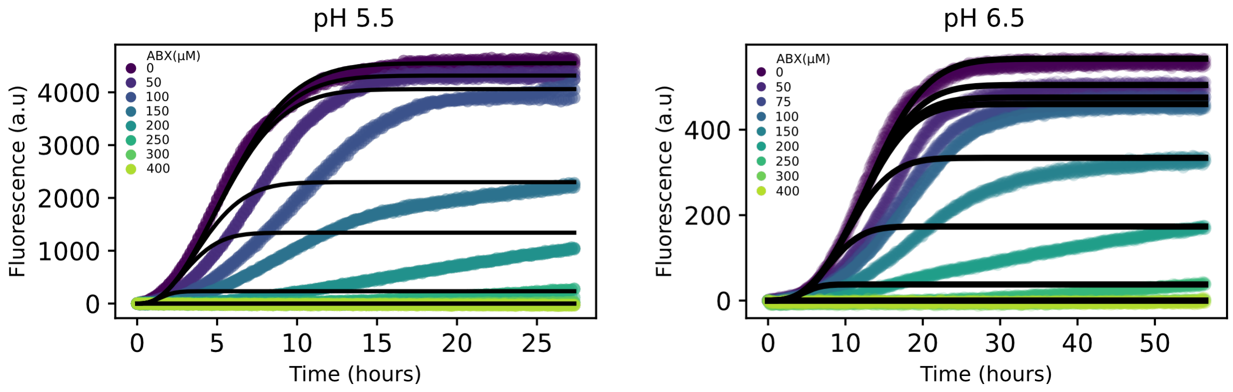


Figure S4: Fitting the availability of DMPS molecules. Fitting the data shown in Fig. 1B and C to a two-step model using the kinetic rates found in the non-inhibited data shown in Fig. 3, while changing the concentration of available DMPS molecules in the system.
